# Product inhibition can accelerate evolution

**DOI:** 10.1101/2022.06.14.496101

**Authors:** Beatrice Ruth, Peter Dittrich

## Abstract

Molecular replicators studied *in-vitro* exhibit product inhibition, typically caused by the hybridization of products into complexes that are not able to replicate. As a result, the replication rate and the selection pressure is reduced, potentially allowing the “survival of everyone”. Here, we introduce a stochastic evolution model of replicating and hybridizing RNA strands to study the effect of product inhibition on evolution. We found that hybridization, though reducing the efficiency of replication, can increase the rate of evolution, measured as fitness gain within a period of time. The positive effect has been observed for a mutation error smaller than half of the error threshold. In this situation, frequency-dependent competition causes an increased diversity that spreads not only within a neutral network but also over various neutral networks through a dynamical modulation of the fitness landscape, resulting in a more effective search for better replicators. The underlying model is inspired by RNA virus replication and the RNA world hypothesis. Further investigations are needed to validate the actual effect of accelerated evolution through product inhibition in those systems.

## Introduction

Replication of polymers is central to the reproduction of organisms and viruses (***Jones et al., 2021***), a key element in major theories of the origin of life (***Gilbert, 1986***), and of interest to make synthetic life (***Adamski et al.,2020***; ***Vay et al., 2019***). Enzyme-controlled (***Haruna and Spiegelman, 1965***) and enzyme-free (***von Kiedrowski, 1986***; ***Zielenski and Orgel, 1987***) (self-)replication has been instantiated *in-vitro* and used to study chemical evolution (***Liu et al., 2020***; ***Salditt et al., 2020***) and to implement bio-molecular procedure (***van Nies et al., 2018***), like the polymerase chain reaction (***Mullis, 1990***).

In *in-vitro* experiments of molecular replication, it has been observed that accumulation of product tends to inhibit the replication process, leading to sub-exponential growth (***von Kiedrowski, 1986***; ***Zielenski and Orgel, 1987***). In fact, product inhibition causes a reduction of the replication rate and reduction of selection pressure, which can lead to the survival of everyone (***Szathmáry, 1991***).

For example, von Kiedrowski (***von Kiedrowski, 1986***) has observed that the concentration of self-replicating hexadeoxynucleotides does not grow exponentially but sub-exponentially. The reason is the formation of hybrids, which act as sinks for single stranded molecules and thus limiting their growth. The effect has been theoretically investigated for replicating RNA sequences by Biebricher et al. (***Biebricher et al., 1985***). They also showed that different single stranded molecules can coexist without cooperative hypercyclic coupling, proposed by Eigen and Schuster (***Eigen and Schuster, 1977***).

Experimentally observations by Rohde et al. (***Rohde et al., 1995***) showed that hybridization leads to a broad mutant distribution of an RNA species replicated by Q*β* replicase. The models introduced in the context of the mentioned studies explain the coexistence of different types of species using rate equations where all molecular types are predefined. Such models do not cover how new types are formed and how mutation influences the time evolution of the population. In contrast our model enables the formation of new types through the simulation of single molecules, here single RNA strands. To avoid the problem of defining all molecular types in advance, our model uses an improved exact stochastic simulation algorithm similar to the Gillespie algorithm (***Gillespie, 1976***) instead of a set of ordinary differential equations.

Although product inhibition supports the “survival of everyone”, an additional directed selection pressure can lead to certain adaptations (***Meszéna and Szathmáry, 2002***) and can cause complex patterns of species formation (***Dittrich and Banzhaf, 2001***). For example, Ito et al. (***Ito and Dieckmann, 2007***) showed that adaptive radiation through intraspecific competition together with weak directional selection of a quantitative trait can lead to rich macroevolutionary patterns involving recurrent adaptive radiations and extinctions. If directional selection is sufficiently weak, evolutionary branching can occur under product inhibition (***Ito and Dieckmann, 2014***). However, because of its narrow scope of evolvability, product inhibition and the resulting parabolic replication has been seen to be of limited relevance for prebiotic evolution (***Szilágyi et al., 2017***), and thus mechanisms circumventing product inhibition have been suggested, like compartments (***Szathmáry and Smith, 1997***) or the formation of intramolecular secondary structures (***Mizuuchi et al., 2020***).

Yet, the quantitative benefit on the fitness gain caused by product inhibition has not been studied in detail (***Szilágyi et al., 2017***). More recent advances have shown that product inhibition not only promotes coexistence but also relaxes the error threshold (***Paczkó et al., 2024***).

For an explicit simulation of RNA sequences evolution Fontana and Schuster (***Fontana and Schuster, 1987***) introduced a fitness function that uses the secondary structure of an RNA sequence to compute its fitness. We follow this approach, because it provides a certain level of realism while the fitness being efficiently computable in polynomial *O*(*n*^3^) time (***Nussinov et al., 1978***). The RNA fitness landscape possesses neutral networks spanning whole sequence space (***Reidys et al., 1997***).

Typical evolution is characterized by quasispecies distributions expanding and drifting on those neutral networks, improving in fitness by contineous and discontineous transitions (***Fontana and Schuster, 1998***; ***Kupczok and Dittrich, 2006***).

RNA viruses profiting from drifting and expanding on neutral networks (***Frost et al., 2018***; ***Lauring, 2020***) may also experience product inhibition during replication in their hosts. The subsequent question than would be if the observed effect of our simulations is transferable to those viruses, contributing to an explanation for their high genotypic variance and quick adaption to new environments (***Frost et al., 2018***; ***Lauring, 2020***).

## Results

### Model

In our model ^1^, a well-stirred population of replicating and hybridizing RNA molecules is simulated. Replication requires a substrate *S* with copy number *N*(*S*), initially set to determine the maximal possible number of RNA molecules. A single stranded RNA molecule with sequence *r* and copy number *N*(*r*) replicates with error by consuming substrate *S* in volume *V* at a rate 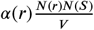, with replication rate constant *α*(*r*). A single stranded RNA molecule decays at a stochastic rate *ϕN*(*r*), here *ϕ* = 1, releasing substrate *S*. A sequence *r* represents that sequence as well as its complement, simplifying the replication model.

Two RNA molecules *r, r*^′^ can hybridize at a rate *β*(*r, r*^′^)*N*(*r*)*N*(*r*^′^)/*V* forming a complex. Note that in case both sequences *r* and *r*^′^ are equal the rate results from *β*(*r, r*^′^)*N*(*r*)(*N*(*r*^′^) − 1)/(2*V*) (***Gillespie, 1976***). Furthermore, we assume that the resulting dimer complex cannot replicate and dissolving of the complex back into two single strands is slow such that it can be omitted. Thus the formed complex can be ignored. To maintain a constant maximal possible number of single stranded RNA molecules the complex is replaced by 2*S*.

In summary, our model consists of the following reactions for RNA sequences *r, r*^′^ ∈ {*A, C, G, U*}^*l*^, which are similar to Epstein’s non-reproductive pairing model (***Epstein, 1979***):

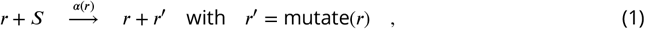

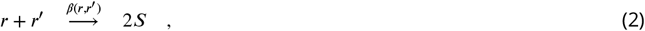

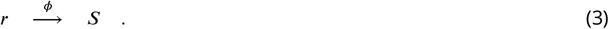

We simulate RNA sequences of fixed length *l*. Typically, we use *l* = 76; additionally, we consider sequences of lengths *l* = 23, 48, 85, and 92. The function mutate(*r*) returns a mutated copy of *r* where each site is mutated with probability *p*. For the replication rate constant *α*(*r*) we take a standard model of RNA evolution, namely the scaled distance *d*_*sec*_(*r, r*_*target*_) of the secondary structure of *r* to a fixed target secondary structure *r*_*target*_; an approach that is said to provide a relatively realistic fitness landscape (***Fontana and Schuster, 1998***): In our case the target secondary structure is the shape of a tRNA (see Materials and Methods). The replication rate constant for a sequence *r* is defined as

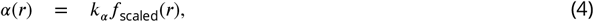

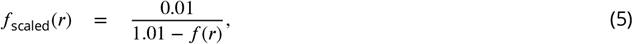

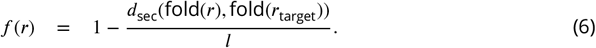

The scaling factor *k*_*α*_ is set to 100. The process of folding a sequence *r* into a secondary structure and the secondary structure alignment *d*_*sec*_(.,.) are computed by the *fold* and *tree_edit_distance* functions of the *ViennaRNA* package (***Lorenz et al., 2011***), respectively.

The hybridization rate constant *β*(*r, r*^′^) is computed from the hybridizing sequences *r, r*^′^ as

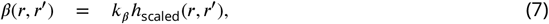

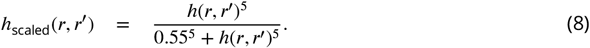

Here, the hybridization strength *k*_*β*_ is varied over the values {0, 0.05, 0.1, 0.3, 1, 3, 4, 10}, where 0 implies no hybridization and increasing values lead to a stronger hybridization influence. The hybridization coefficient *h*(*r, r*^′^) depends on the hybridizing single strands *r* and *r*^′^. Roughly, the more similar they are the more likely they hybridize. Because we assume RNA sequences with a fixed length *l*, we can compute the hybridization coefficient *h*(*r, r*^′^) from the sum of Gibbs free energy contribution of each base pair considering the adjacent base pairs (see Methods for details). Note that this leads to a more realistic model than using the Hamming distance, because the Watson-Crick base pairs CG UT have different contributions as they allow a different number of hydrogen bonds. Further note that our conclusions also hold for alternative scaling functions (Appendix 3).

For simulation, we usually generate a random initial population containing *N*(*r*) = 50 copies of a randomly generated RNA sequences *r* of length *l* = 76 and *N*(*S*) = 9950 substrate units, allowing a total population of *n* = 10000 RNA sequences. We let the population evolve by an improved exact stochastic simulation algorithm similar to the Gillespie algorithm (for details and a proof of correctness see Appendix 2). The new algorithm is necessary because the standard Gillespie algorithm cannot cope with the large number of possible hybridization reactions, which scale quadratically with *n*.

Time *t* is measured in generations. In one generation, one new RNA sequence per sequence in the population is generated by replication. Technically, *t* is incremented by one fraction of the current population size, i.e. *t* ← *t* + 1/*N*(*r*), in each replication event (see Appendix 1).

### Product inhibition by hybridization can accelerate evolution

We measure the rate of evolution by the fitness *f*_scaled_ of the population’s best sequence after time Δ*t*. Figure 1 shows the reached fitness after Δ*t* = 100, 300, and 1000 generations, respectively. We have omitted the transient phase from 0 to generation 100, because the behavior is strongly dependent on our choice how to initialize the population. During this transient phase, the amount of RNA sequences typically grows quickly (within 10 generations) from 50 towards the maximum carrying capacity of 10000. Furthermore, the diversity grows due to mutations and hybridization. Because initial diversity is low (all sequences are equal at Δ*t* = 0), hybridization is likely, and thus a system without hybridization gains fitness faster during this transient phase (Appendix 3 – figure 1A).

**Figure 1.**
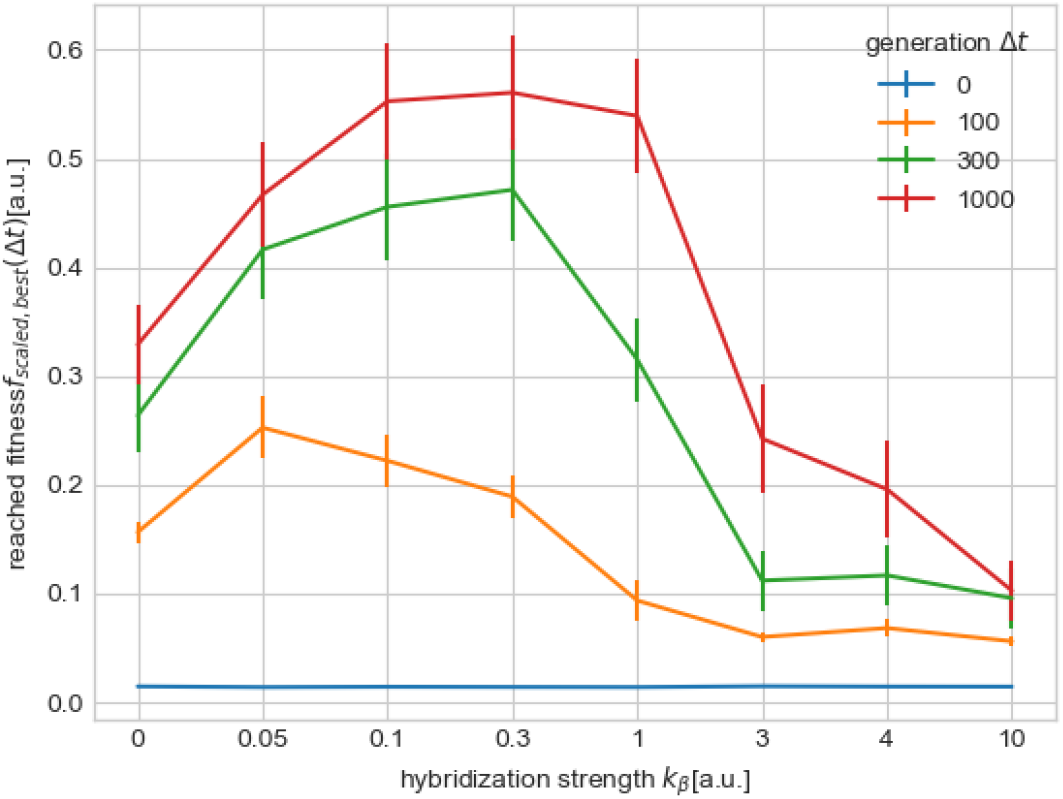
Effect of hybridization on the efficiency of evolution at a moderate mutation rate *p* = 0.01 per base. Evolution rate is measured in reached fitness *f*_scaled,best_ of the best individual under the presence of hybridization *k*_*β*_ > 0 compared to no hybridization *k*_*β*_ = 0. Even in early generations Δ*t* = 100 (orange dots) the evolved fitness in populations with a low hybridization rate *k*_*β*_ <= 1 is at a comparable height to that of populations without hybridization. Late generations Δ*t* > 300 (green and red dots) reveal not only an accelerated evolution for populations with hybridization 0 < *k*_*β*_ <= 1 but also a shift from the highest overall achieved fitness from *k*_*β*_ = 0.05 at generation Δ*t* = 100 to *k*_*β*_ = {0.1, 0.3} at generation Δ*t* = 1000. The fitness *f*_scaled,best_ of the best sequence of a population is averaged over 50 simulations, with error bars showing the standard error of the mean, using volume *V* = 1.

Interestingly, as time progresses (Δ*t* > 100), a population with hybridization, causing product inhibition, can reach a significantly higher fitness than the same population without hybridization. This effect is maintained over many generations (Δ*t* = 1000), resulting in a widening gap in fitness reached between populations with and without hybridization. The higher rate of evolution becomes visible at medium hybridization strengths. In particular, the hybridization factor *k*_*β*_ can vary over one magnitude of range, that is, from *k*_*β*_ ≈ 0.05 to ≈ 0.3 (Figure 1).

In order to rule out effects resulting from the chosen secondary structure as target, we have tested five additional target structures from sequences of length 23, 48, 86, 90 and 91 bases, respectively. Results are shown in Figure 2 and Appendix 3 – figure 1. The effects of low hybridization rates causing strong fitness gain in early generations and of medium hybridization rates causing an increased fitness gain in later generations are in line with the previous target structure shown in Figure 1. Thus, the choice of a target structure seems to have only a minor impact on the overall observed accelerated evolution under product inhibition.

**Figure 2.**
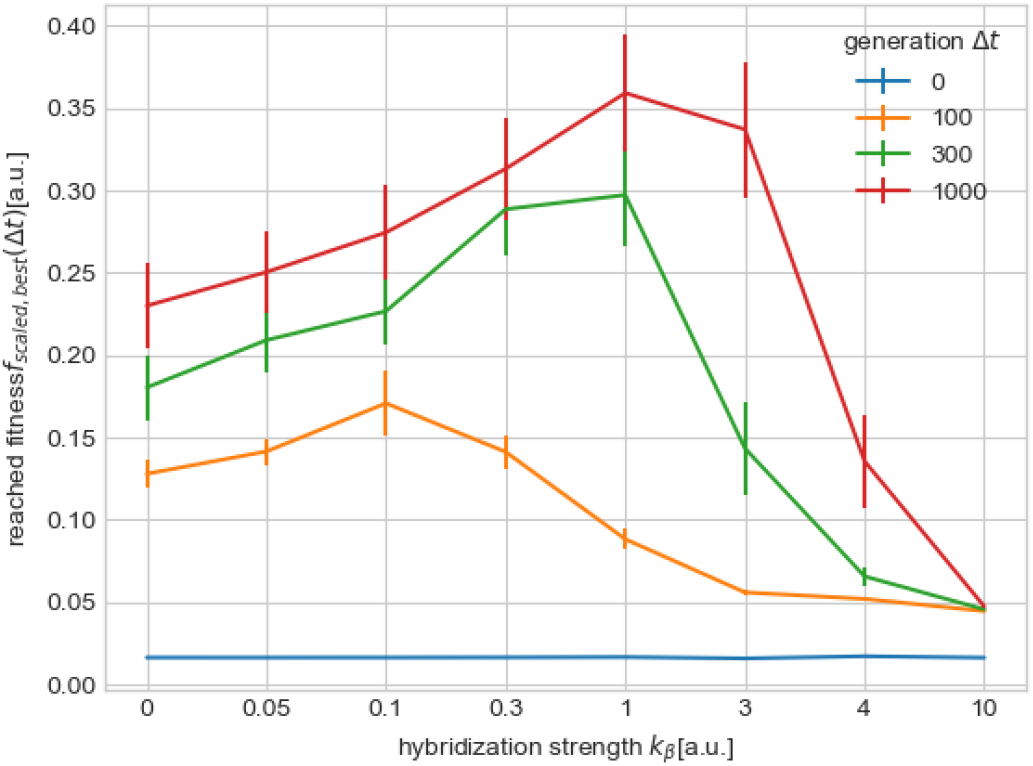
Effect of hybridization on the efficiency of evolution at a moderate mutation rate (*p* = 0.01) with Homo sapiens Small nucleolar RNA U3 (Appendix 3, Tab. 1). The fitness *f*_scaled,best_ of the best sequence of a population is averaged over 50 simulations, with error bars showing the standard error of the mean, using volume *V* = 10. Rising generations not only reveal a stronger difference in reached fitness but also the range of the hybridization rate associated with accelerated evolution widened till *k*_*β*_ = 3.

### Hybridization inhibits evolution when mutation is almost at the error threshold

The error threshold *p*_*err*_ is a limit of the mutation probability of a base above which mutation will destroy the sequence information over time (***Eigen, 1971***; ***Eigen and Schuster, 1977***). In our model, without hybridization (*k*_*β*_ = 0) we have *p*_*err*_ ≈ 0.025 per base, in line with ***Kupczok and Dittrich (2006***), Figure 3.

**Figure 3.**
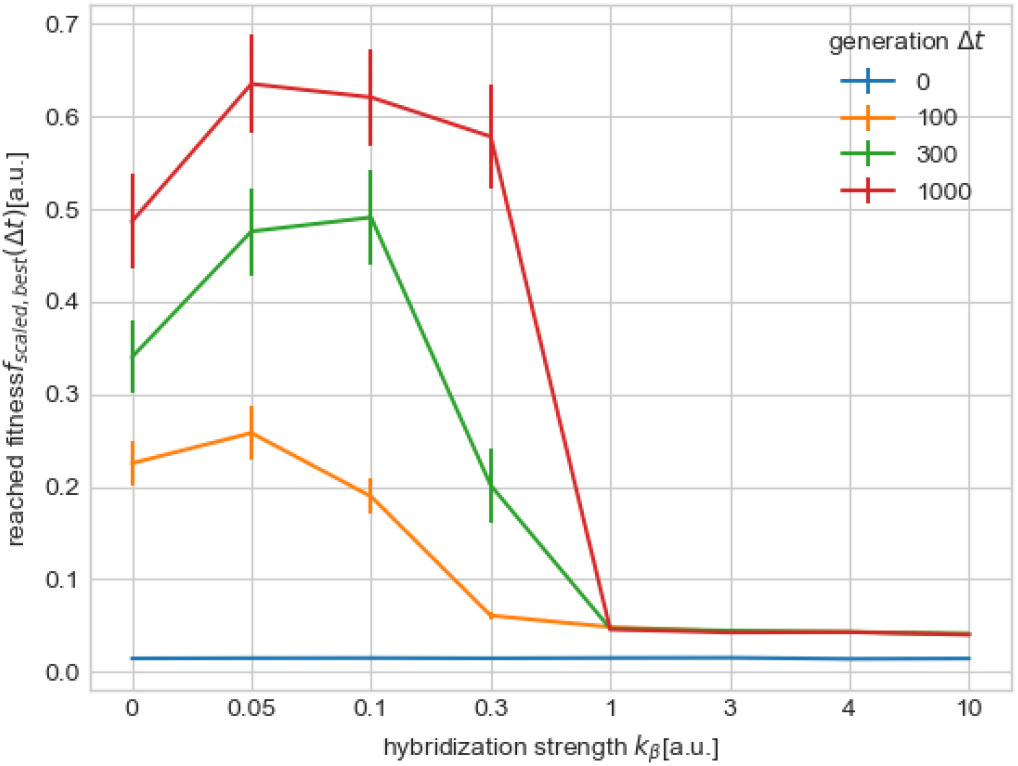
Effect of hybridization on the efficiency of evolution at a high mutation rate *p* = 0.02, close to the error threshold (*p*_*err*_ ≈ 0.02). Early Δ*t* = 100 (orange dots) and late Δ*t* = {300, 1000} (green and red dots) generations reveal upon increasing hybridization *k*_*β*_ a decrease in efficiency of evolution. For high hybridization *k*_*β*_ >= 1 even no fitness improvement is observable. Only low hybridization rates *k*_*β*_ = {0.05, 0.1} are able of reaching similar values for the fitness in early generations. Surpassing *k*_*β*_ = 0 in late generations is even achieved for mildly increased *k*_*β*_ <= 0.3. Mean and standard error of the mean shown, based on 50 simulations each, using volume *V* = 1.

Simulations performed with a high mutation rate of *p* = 0.02 showed that if the mutation probability is close to the error threshold, hybridization can have a negative effect on the rate of evolution (Figure 3). For example, at a moderate hybridization strength *k*_*β*_ = 1, which leads to a higher rate of evolution at a moderate mutation rate *p* = 0.01 (Figure 1), evolution is fully inhibited at a high mutation rate close to the error threshold (Figure 3). This is in line with the fact that hybridization reduces the overall growth rate of a sequence and causes a bifurcation from a stable to an unstable regime. We can also see in Figure 3 that with increasing hybridization strength, the effective error threshold decreases.

### Decreasing population density can accelerate evolution under hybridization

As high mutation rates close to the error threshold have a negative effect on the fitness development of populations with hybridization, moderate mutation rates reveal a more benefiting evolution for populations with hybridization compared to populations without hybridization (Figure 1). For such a moderate mutation rate, there is a regime of hybridization strengths (1 ≳ *k*_*β*_) where increasing the volume leads to an increased rate of evolution (Figure 4). Increasing the volume is equivalent to decreasing the population density by keeping the number of sequences and substrate constant. Leading to a decrease of replication and hybridization rates while the first order decay rate stays unchanged.

**Figure 4.**
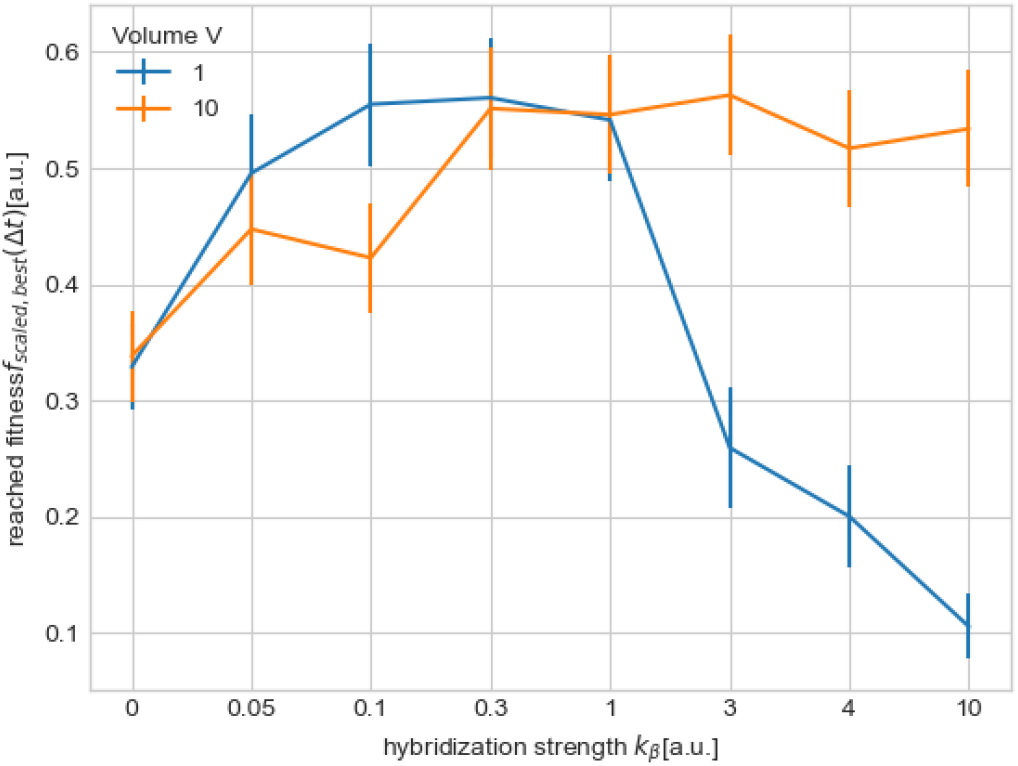
Influence of the volume on the efficiency of evolution. An increase in volume to *V* = 10 (orange dots) leads to an decreased fitness *f*_scaled,best_ for small hybridization rates (*k*_*β*_ ≤ 0.1). However, for large hybridization rates (*k*_*β*_ > 1) it has a positive effect. Note that a decrease in volume is equivalent to an increase of the second order reaction rates (replication and hybridization rates, here). Mean and standard error of the mean shown, based on 25 simulations each, using volume *V* = 10, a moderate mutation rate *p* = 0.01, and Δ*t* = 1000.

### Improvement by hybridization could be explained by a broader mutant spectrum

We measure population diversity and structure by Hamming distance among pairs of sequences. With increasing hybridization strength from *k*_*β*_ = 0 to *k*_*β*_ = 10 the mean Hamming distance increases roughly linearly from 12 to 40 bases (Figure 5). Note that for *k*_*β*_ = 10 the mean Hamming distance is larger than the mean distance between two random sequences (*l*/2 = 38). Further note that at such a high hybridization strength there is no evolutionary progress anymore (Figures 1-4).

**Figure 5.**
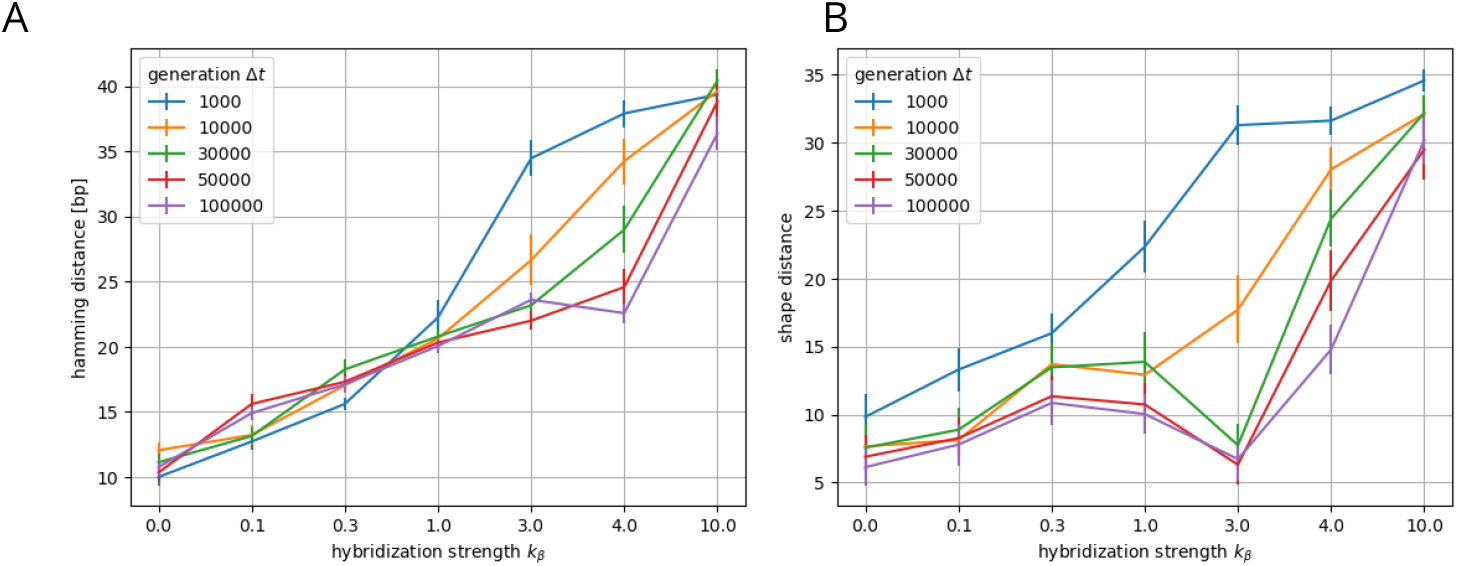
Average hamming distance (A) and average shape distance (B) between two sequences in a population depending on the hybridization strength *k*_*β*_ and generation Δ*t*. The pairwise hamming distance in small generations Δ*t* = {1000, 10000} increases dependent on a growing hybridization strength *k*_*β*_, which is inline with the pairwise distance of their secondary structures, shapes. Where hamming distances in later generations Δ*t* = {30000, 50000, 100000} have an reduced increase based on a growing hybridization strength, their respective secondary structures have a minimal distance at hybridization strength *k*_*β*_ = 3.0. Mean and standard error of the mean shown, each based on 25 simulations with a population size of 100 sequences, using volume *V* = 10, and a moderate mutation rate *p* = 0.01 on a much smaller population of *N* = 100 sequences.

So, an improved rate of evolution through hybridization coincides with a moderately broader mutant spectrum. Despite the absence of an increased number of clusters in sequence space (Figure 6), the hybridizing population seems to explore the sequence space more effectively, owing to its broader mutant spectrum (Figure 5). While a population without hybridization usually occupies one neutral network during its exploratory phase (***Fontana and Schuster, 1998***), a population with hybridization might occupy stably more than one neutral network (cf. Figure 7 and (***Dittrich and Banzhaf, 2001***)).

**Figure 6.**
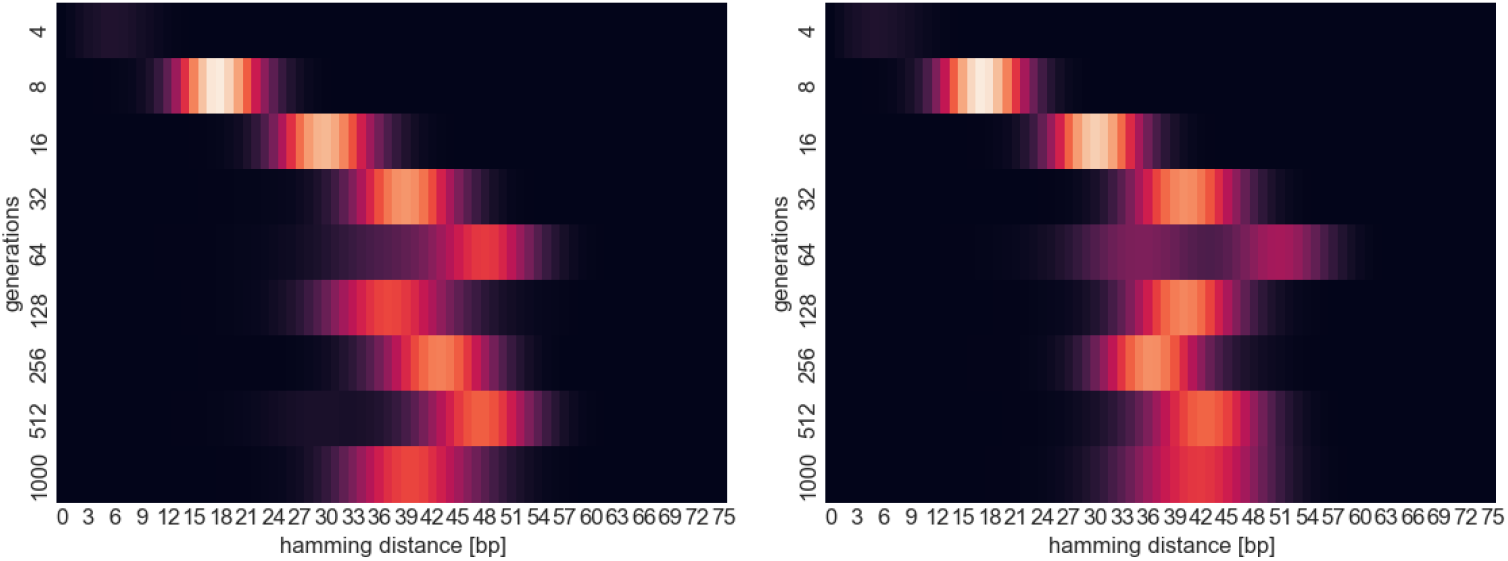
Comparison of the temporal progression of the hamming distance distribution between a population without (left) and a population with hybridization (*k*_*β*_ = 3, right) considering a volume *V* = 1 and a moderate mutation rate *p* = 0.01.

**Figure 7.**
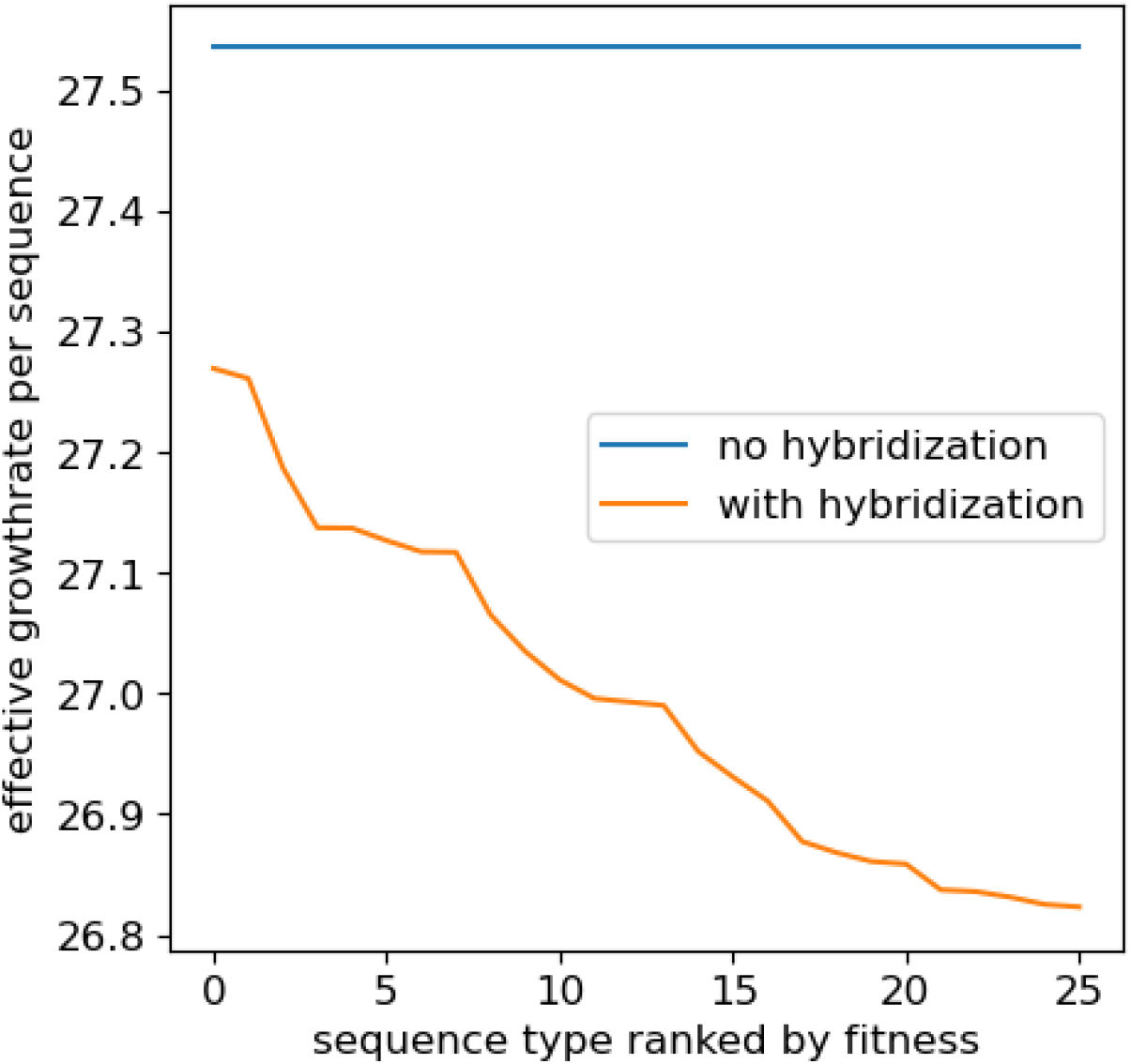
Modulation of the effective fitness landscape (effective growth rate) due to hybridization. Neutral networks vanish under the presence of hybridization events as there are no longer different sequence types with the same effective growth rate. Depicted data shows the first 26 sequence types ranked by the effective growth rate in the final populations obtained in Figure 6, with volume *V* = 10, mutation rate *p* = 0.01 and *k*_*β*_ ∈ {0, 3}.

### Hybridization dynamically modulates the fitness landscape

Here, we define the effective fitness of a sequence *r* with respect to a population *P* as the effective growth rate of that sequence. This rate is not only determined by the sequence’s replication and decay rate but also includes the negative effect caused by hybridization with other sequences of the population. Thus, the evolving population dynamically modulates its effective fitness landscape through hybridization, cf. (***Dittrich and Banzhaf, 2001***). In a population without hybridization, the first 26 sequence types show the same effective growth rate. In contrast, a population with hybridization each sequence type has, in general, a different effective growth rate (Figure 7). As a consequence, there are no neutral networks anymore because the change in number of one sequence *r* leads to an individual change in effective fitness in all sequences, depending on their general probability of hybridizing with that sequence *r*.

### Hybridization adds an additional selection pressure on a sequence’s GC content

Figure 8 shows that with increasing hybridization strength the average GC content of the sequences decreases. This effect can be explained by the difference of energy contributions between GC and AU base pairs, which are the basis for the rate of a hybridization reaction. The observed selection pressure towards lower GC-content in order to avoid hybridization increases with rising chances of a hybridization event but scales down with increasing volume making it harder for sequences to collide. It has also been observed in a simulation model by ***Paczkó et al. (2024***).

**Figure 8.**
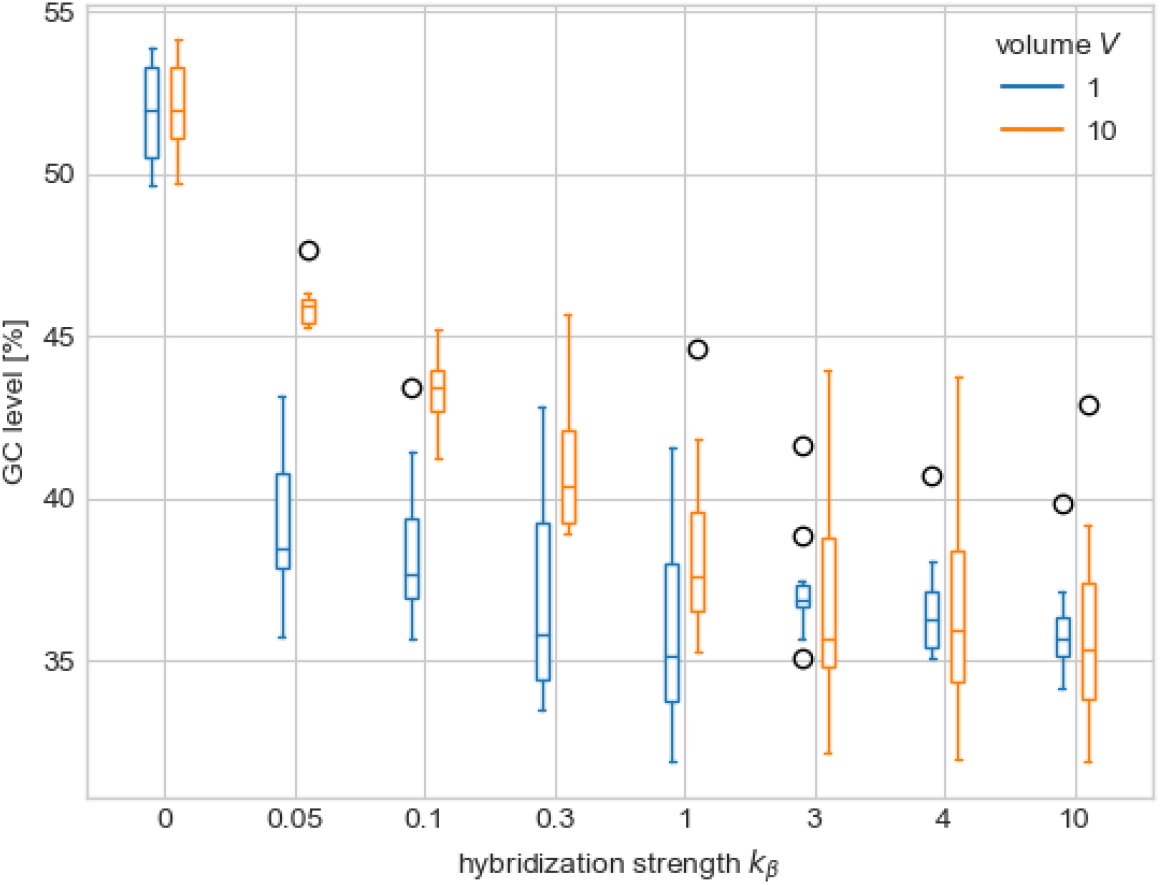
Influence of hybridization on the GC content depending selection of sequences. An increased hybridization strength increases the selection pressure on the GC content, leading to a decrease in GC level. Depicted data considers the range of GC level of the sequence with highest fitness of each simulation, with volume *V* = {1, 10} and mutation rate *p* = 0.01, from generation Δ*t* = 100 up to generation Δ*t* = 1000. Coloured boxes contain quartiles 1 to 3 of the range of the GC level with their whiskers having 1.5 times the length of the box.

To rule out that the implicit selection pressure towards low GC-content is causing the increased rate of evolution through product inhibition, we performed “unweighted” simulations where GC and AU base pairings are treated equally in calculating hybridization rates. In these simulation no implicit selection pressure on the GC-content is created through hybridization. Figure 9 shows that neglecting GC-content during hybridization leads to an even more accelerated evolution in hybridizing populations. Starting around generation 300, a significant difference emerges between the fitness levels reached with weighted and unweighted hybridization. This indicates that the acceleration of evolution is primarily driven by product inhibition through hybridization and not by the implicit selection pressure on the GC-content. Additional experiments using variations of the hybridization model confirmed this beneficial effect on the speed of evolution (Appendix 3 – figure 3). Finally, note that the sequence of our target structure has a high GC-content. So, selection pressure towards low GC-content does not push evolution towards this sequence.

**Figure 9.**
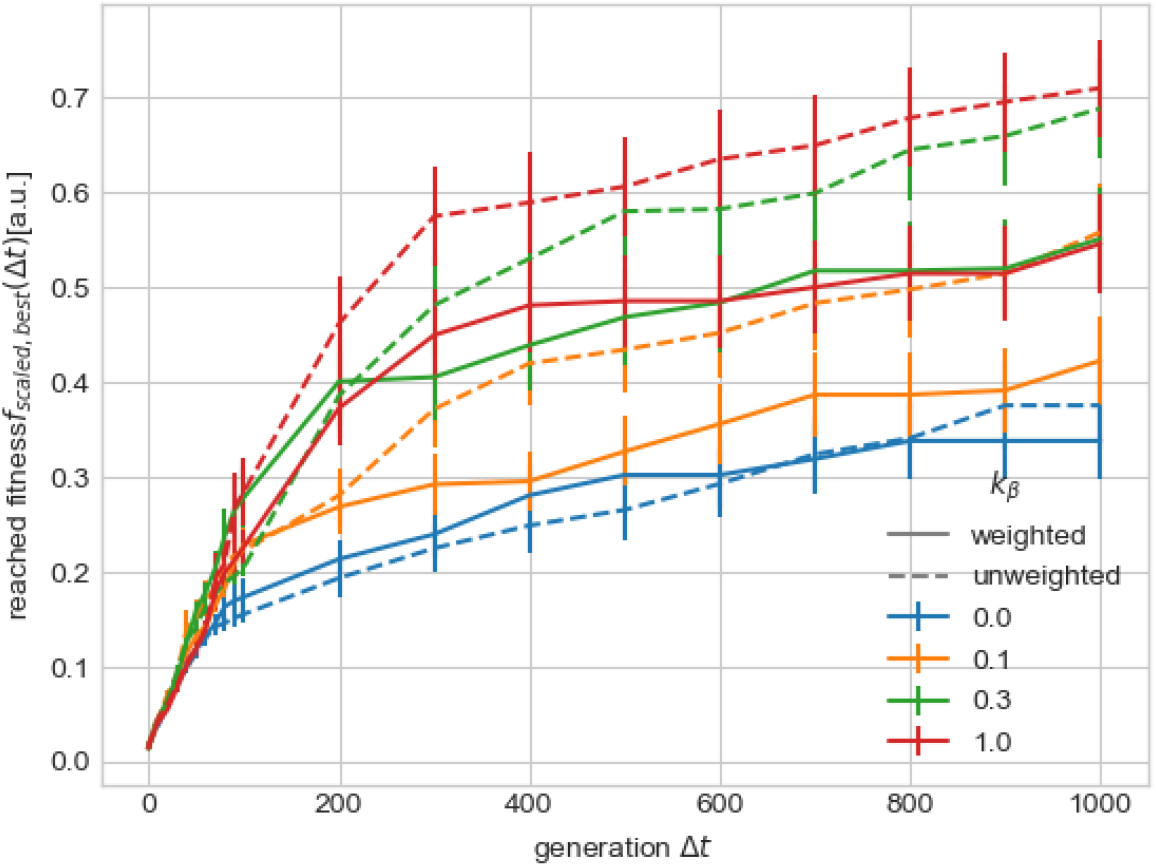
Comparison of hybridization with a homologous, unweighted, hybridization (dashed line) and a weighted hybridization (solid line) respecting the differences of the individual base pairings. If the effect of the individual base pairings is neglected an even higher fitness is reachable for the presented hybridization strengths *k*_*β*_ ∈ {0.1, 0.3, 1.0}. The acceleration of evolution is greater if hybridization is treated unweighted for the individual base pairings, suggesting that a pressure on the GC content does not lead to an accelerated evolution. Mean and standard error of the mean shown, based on 50 simulations each, using volume *V* = 1, and a moderate mutation rate *p* = 0.01.

## Discussion

A novel and proven exact extension of the Gillespie stochastic simulation approach allowed us to perform exact stochastic simulations of the evolution of replicating and hybridizing RNA sequences. Simulations showed that hybridization, though reducing the rate of replication, can increase the rate of evolution, measured as fitness gain within a period of time. We have measured time in generations, which is reasonable because we assumed that replication requires a resource available at a limited rate. This limited availability of the resource implied a maximal carrying capacity of the population and thus competition for the resource, causing selection.

The positive effect of hybridization has been observed clearly for a mutation rate *p* that has a certain distance to the error-threshold, e.g., half of the error-threshold mutation rate. For such a “safe” mutation rate, hybridization has led to an improvement in fitness over a wide range of hybridization strengths and evolution times, e.g., 100 to 1000 generations. When the population replicates at the error threshold, product inhibition can easily push the population over this threshold, causing a loss of sequence information. Therefore, we expect that in a system that utilises product inhibition to accelerate evolution, the mutation rate is at a certain distance from the error threshold.

Positive effects on the evolution rate have been observed for hybridization parameters ranging from *β*_*k*_ = 0.05 to *β*_*k*_ = 1, or more for diluted systems with volume *V* = 10 (Figure 4). This range corresponds to relative amounts of inhibited products (dimers) to uninhibited sequences (single strands) between 18% to 45% (Appendix 3 – figure 2A) and 7% to 35% (Appendix 3 – figure 2B). Thus, the positive effect could be expected even in systems with relatively high amplification efficiency, like Q*β* replicase under “ideal” conditions, which we would align to a *β*_*k*_ ≈ 0.05. Note that other RNA-dependent RNA polymerases display much stronger product inhibition, suggesting that evolution can, at least in principle, move the rate of product inhibition into a beneficial regime by adapting the replicase system.

We explained the positive effect of hybridization by an increase in diversity. With hybridization populations are able to expand more freely in sequence space, providing more opportunities for a benefiting mutation in terms of a higher replication rate. We note that the effect is closely related to the limiting similarity principle (***MacArthur and Levins, 1967***), saying that phenotype difference between two species on the scale of the competition width is required for coexistence (***Szabó and Meszéna, 2006***).

The evolutionary phenomenon investigated in this study could be relevant and further explored in: prebiotic evolution, where replicating polymers hypothetically emerged that were very likely subject to product inhibition; in biotic evolution, e.g., where RNA strands of viruses replicate within a biological cell; or in artificial molecular/chemical evolution, where product inhibition might be used to evolve molecules with desired properties more efficiently.

### History

A first version of this paper has been published in bioRxiv (***Ruth and Dittrich, 2022***). Here, for the new version, we extended the simulation experiments to larger populations and performed simulations for five additional target structures, supporting the original findings with more data.

## Methods and Materials

### Hybridization Coefficient *h*(*r, r*^′^)

For a more realistic model of hybridization probability upon colliding sequences *r* and *r*^′^ instead of pure sequence similarity the change of Gibbs free energy upon dimer formation based on nearest-neighbor thermodynamics of oligonucleotides is used (***SantaLucia, 1998***).

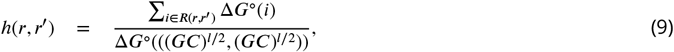

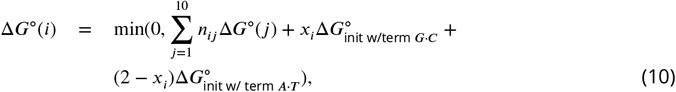

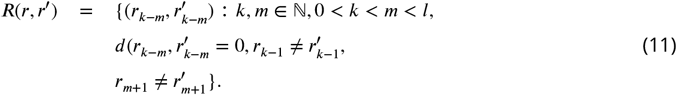

The hybridization coefficient *h*(*r, r*^′^) itself is thereby the normalized sum of the change of Gibbs free energy of the sequence pair (*r, r*^′^) compared to the maximal change of Gibbs free energy via the sequence pair ((*GC*)^*l*/2^, (*GC*^*l*/2^)) with *l* being the length of the sequences *r* and *r*^′^. Only matching regions *R*(*r, r*^′^) contribute to hybridization through the possible formation of hydrogen bonds. As only substitution mutations take place the chance of improving matching regions by insertion of gaps is very small such that it can be omitted. The change of Gibbs free energy in one region Δ*G*^°^(*i*) is dependent on the number of flanking *G* · *C* pairs *x*_*i*_ and the numbers of possible base sequences *n*_*ij*_ with their change of free energy parameter is taken correspondingly from Table 1 (***SantaLucia, 1998***).

**Table 1.** *Unified NN-parameters* as in (***SantaLucia, 1998***). Parameter is the energy difference of the given basepair combination upon hybridization. Basepair combination VW/XY corresponds to sequence sections VW and YX in 3^′^-to-5^′^ orientation. With respect to RNA T is substituted by U for the present model.

| j | sequence | parameter,<br>kcal/mol |
| --- | --- | --- |
| 1 | AA/TT | -1.00 |
| 2 | AT/TA | -0.88 |
| 3 | TA/AT | -0.58 |
| 4 | CA/GT | -1.45 |
| 5 | GT/CA | -1.44 |
| 6 | CT/GA | -1.28 |
| 7 | GA/CT | -1.30 |
| 8 | CG/GC | -2.17 |
| 9 | GC/CG | -2.24 |
| 10 | CC/GG | -1.84 |
| | Init. w/term. $G \cdot C$ | 0.98 |
| | Init. w/term. $A \cdot T$ | 1.03 |

### Target Structure

Like Fontana (***Fontana and Schuster, 1998***) and Kupczok&Dittrich (***Kupczok and Dittrich, 2006***) we used a shape of a tRNA as the target secondary structure obtained from the sequence *r*_*target*_ = ε*GGGCAGAUAGGGCGUGUGAUAGCCCAUAGCGAACCCCCCGCUGAG CUUGUGCGACGUUUGUGCACCCUGUCCCGCU*ε

giving

*f old*(*r*_*target*_) = ((((((…((((……..)))).(((((…….)))))…..((((.((….)))))).))))))…..

The other sequences we used can be found in Appendix 3 3 1.

### Stochastic Simulation

The new algorithm to get stochastic correct trajectories is derived from the Gillespie algorithm (***Gillespie, 1976***) and can be found in the appendix 1 1. The major difference arises from the separation of selecting the reaction type (replication, hybridzation, decay), first, and then selecting the actual reaction with its educts (***Rosenberger et al., 2021***). This allows the formulation of implicit chemical reactions such that not every sequence needs a separate set of replication, hybridization, and decay reactions. The following lemma ensures the correctness of the new algorithm.

#### Lemma 1.

*The developed algorithm is equivalent to the Gillespie algorithm by generating a statistically correct trajectory*.

*Proof*. Both algorithms are equivalent if the following criteria are met:

1. The probability of choosing a certain sequence is the same in both variants.
2. The probability of choosing a certain reaction is the same in both variants.
3. The time interval between two successive reactions is the same in both variants.

These three criteria are proven in the Appendix 2.

## Data availability

The data for this paper has been generated by simulation of the model introduced in this paper and using the target sequences listed in Appendix 3. The code available at https://git.uni-jena.de/ne78xoy/hr-sim-newgillespie, to be archived at Zenodo.

## Acknowledgments

We acknowledge funding by the *Deutsche Forschungsgemeinschaft* (Project 419138358) and by the Ministry for Economics, Sciences and Digital Society of Thuringia (TMWWDG), under the framework of the Landesprogramm ProDigital (DigLeben-5575/10-9).

## Appendix 1

### Simulation algorithm

#### Initialization

(a) Set the time variable *t* = 0.
(b) Specify and store the implicit molecule categories *S*_sequence_ and *S*_substrate_ with their numbers *n*_sequence_ and *n*_substrate_.
(c) Specify and store the split into the explicit molecule categories *S*_sequence,*i*_ and *S*_substrate,*s*_ with their numbers 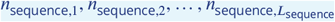 and *n*_substrate,*s*_(In this case: *n*_substrate,*s*_ = 100). The total number of sequences is computable by 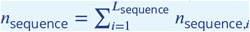.
(d) Specify and store the implicit chemical reactions *R*_*μ*_ with *μ* ∈ {*replication, hybridization, decay*} as *S*_sequence_ + *S*_substrate_ → 2*S*_sequence_ (replication), 2*S*_sequence_ → 2*S*_substrate_ (hybridization) and *S*_sequence_ → *S*_substrate_ (decay).
(e) Calculate and store the quantities *c*_replication_ = *k*_replication_/*V*, *c*_hybridization_ = *k*_hybridization_/*V*, *c*_decay_ = *k*_decay_ (*k*_*μ*_ corresponds to the rate constant and *V* is the volume).
(f) Specify and store the calculation rules 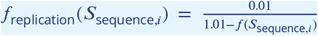 with *f* (*S*_sequence,*i*_) representing the fitness of the explicit sequence *i*, 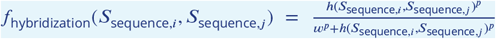 with *h*(*S*_sequence,*i*_, *S* _sequence,*j*_) representing the hybridization coefficient of the explicit sequences *i* and *j, w* and *p* correspond to the turning and exponent of the sigmoid function, and *f*_*decay*_(*S*_*sequence,i*_) = 1 to model the explicit chemical reactions.
(g) Store the maximal achievable values of those calculation rules *e*_*replication*_ = 1, *e*_*hybridization*_ = 1 and *e*_*decay*_ = 1.
(h) Calculate and store all 3 reaction propensities 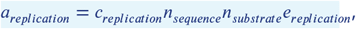 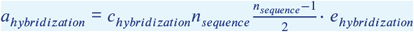 and *a*_*decay*_ = *c*_*decay*_ *n* _*sequence*_.
(i) Specify and store a series of sampling times *t*_1_ < *t*_2_ < …, and also a stopping time *t*_*stop*_. (***Gillespie, 1976***)

#### Timestep

Generate a random number *r*_1_ uniformly distributed on [0, 1] to determine the time interval 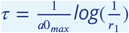 with *a*0 = *a*_*replication*_ + *a* _*hybridization*_ + *a*_*decay*_.

#### Reaction type

(a) Calculate the contribution of *a*_replication_, *a*_hybridization_ and *a*_decay_ to their sum as 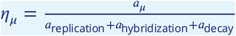 with replication hybridization decay.
(b) Proportional selection of the reaction type *R*_*μ*_ based on *η*_*μ*_.

#### Reaction probability

(a) Proportional selection of the required sequence/s based on their relative frequency 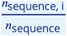.
(b) Calculate the probability *p*_*μ*_ for the chosen reaction type *R*_*μ*_ taking into account the selected sequence/s 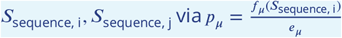 if *μ* ∈ {*replication, decay*} else 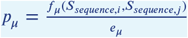.

#### Update

(a) Advance time *t* by the random generated time step *t* = *t* + *τ* in step 2 time step.
(b) Generate a random number *r*_2_ uniformly distributed on [0, 1]. If *r*_2_ < *p* remove all educts and add all products of *R*_*μ*_ else proceed with the next step iteration.
(c) If replication takes place add once the removed sequence and generate a mutated sequence from the explicit sequence. If the mutated sequence is already present add 1 to *n*_*sequence,i*_ of the corresponding sequence *S*_*sequence,i*_ else *L*_*sequence*_ = *L*_*sequence*_ + 1 and 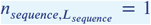 with 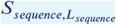 being the newly mutated sequence.
(d) Delete all *S*_*sequence,i*_ with *n*_*sequence,i*_ = 0. For each deleted *S*_*sequence,i*_ adjust the following indices *i* + *x* with *x* ∈ ℕ * by *i* + *x* − 1 and *L*_*sequence*_ = *L*_*sequence*_ − 1.
(e) Recalculate 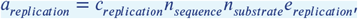 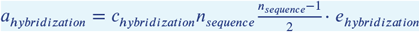 and *a*_*decay*_ = *c*_*decay*_ *n*_*sequence*._

#### Iteration

(a) If *t* has just been advanced through one of the sampling times *t*_*i*_, read out the current molecular population values 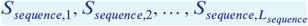.
(b) If *t* > *t*_*stop*_, or if no more reactions are possible (all *a*_*μ*_ = 0), terminate the calculation; otherwise, return to Step 2 time step.

## Appendix 2

### Proof of Lemma 1

For proving Lemma 1 from the main text

#### Lemma 2.

we need to prove the following three sub-lemmata:

#### Lemma 2.1.

*The probability of choosing a certain sequence is the same in both variants. Proof*. Through the rate equations the following is obtained for the Gillespie algorithm:

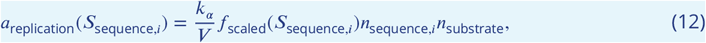

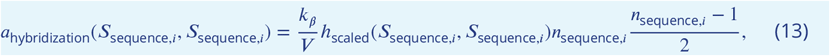

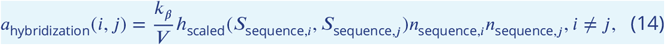

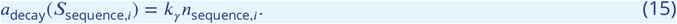

And for the developed algorithm:

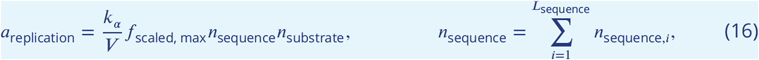

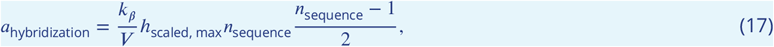

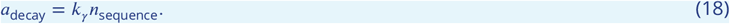

in the developed algorithm only after choosing the reaction type the explicit molecules in this case sequences are randomly chosen based on their relative abundancies reacting accordingly to their specific reaction parameter (*f*_scaled_(*S*_sequence,*i*_), *h*_scaled_(*S*_sequence,*i*_, *S*_sequence,*j*_)), leading to:

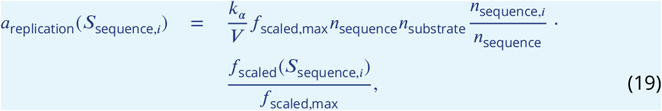

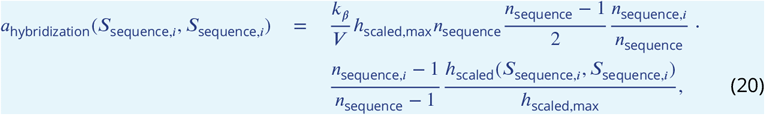

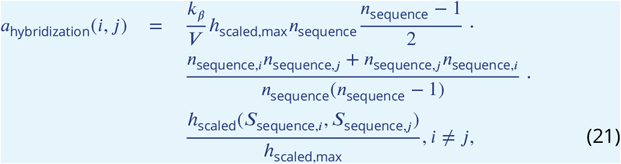

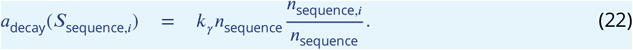

Since *n*_sequence_, *f*_scaled,max_ and *h*_scaled,max_ in the equations above cancel each other out the equations of the Gillespie algorithm are obtained. Thus leading to the conclusion that in both cases a certain sequence has the same probability to undergo a reaction.

#### Lemma 2.2.

*The probability of choosing a certain reaction is the same in both variants. Proof*. The selection of a reaction type in the Gillespie algorithm:

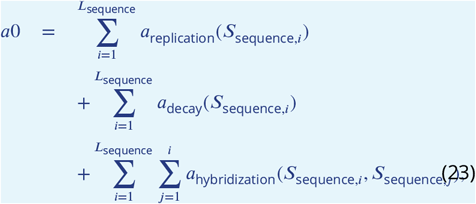

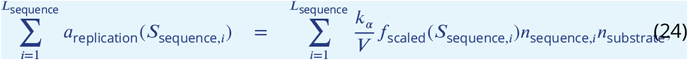

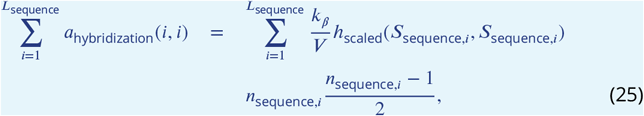

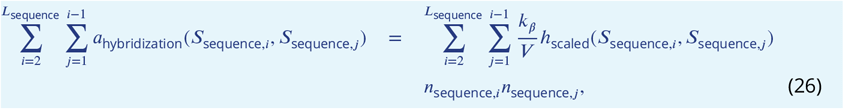

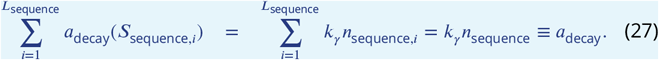

On the other hand the selection of a reaction type in the developed algorithm:

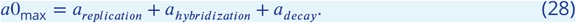

Reshaping the equation of the replication sum in the Gillespie algorithm:

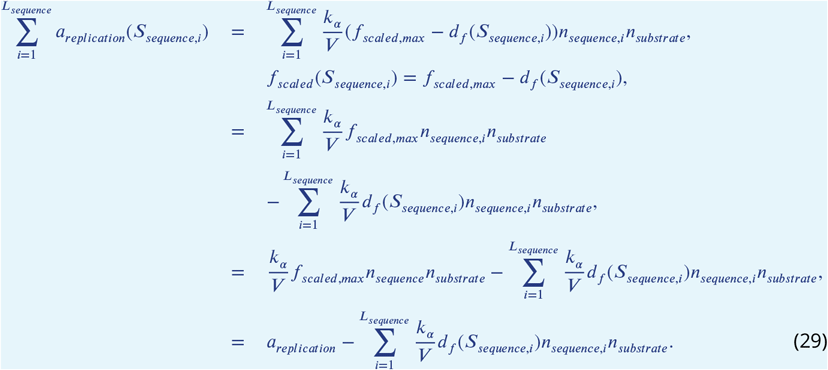

Respecting the fact of denying a replication in the developed algorithm:

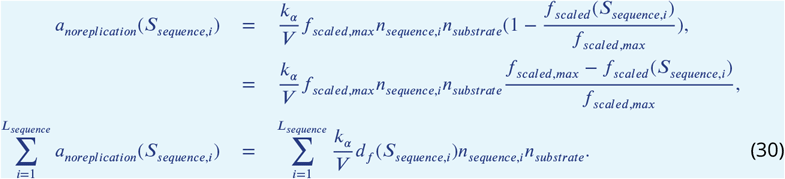

Since the obtained equation is equal to the difference of the direct reaction selection between the two algorithms both algorithms posses the same possibility for a replication event.

Reshaping the hybridization sum for equal sequences of the Gillespie algorithm:

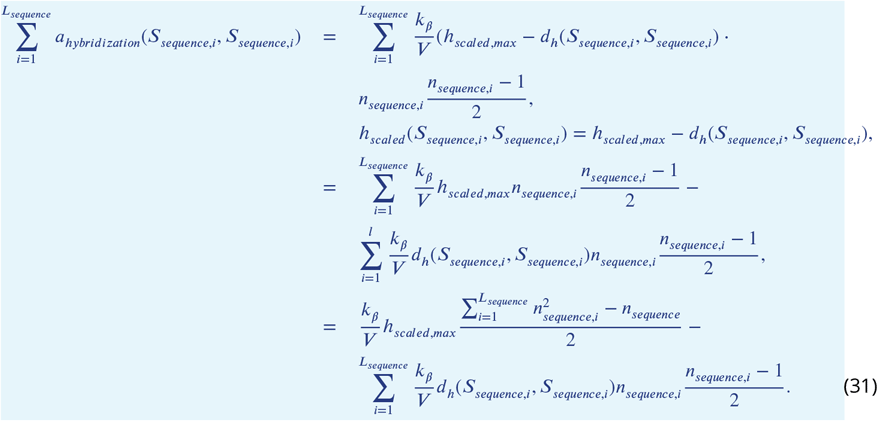

Reshaping the hybridization sum for unequal sequences of the Gillespie algorithm:

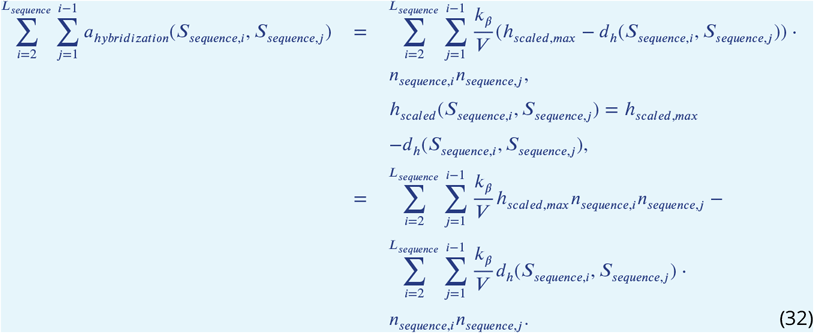

A simplified view via adding the terms with *h*_*scaled,max*_ of hybridization with equal and unequal sequences leads to:

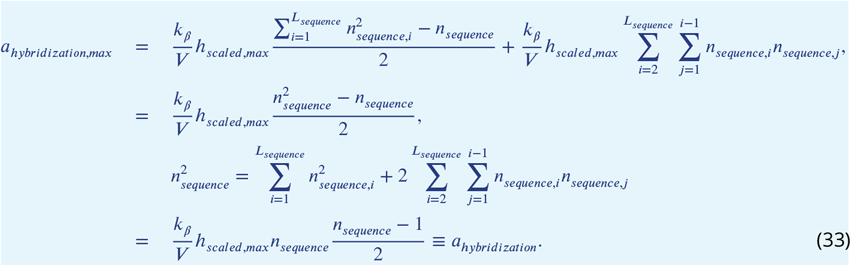

The difference between both algorithms immediately after selection of hybridization is the sum of the terms containing *d*_*h*_(*S*_*sequence,i*_, *S*_*sequence,j*_):

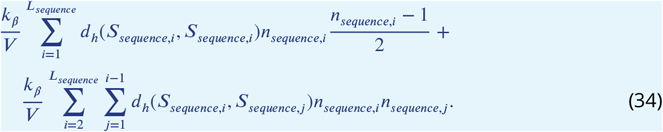

Respecting the fact of denying a hybridization after the selection of the explicit sequences in the developed algorithm with probability 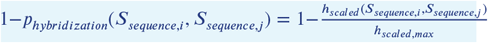.

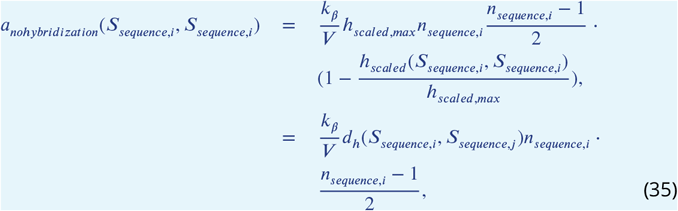

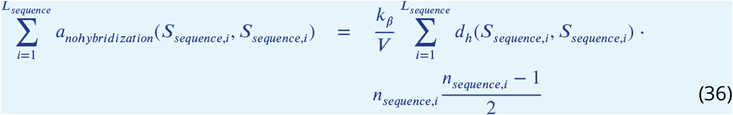

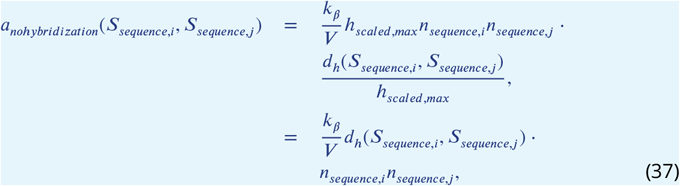

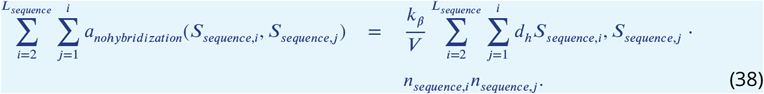

The possibility of denying a hybridization event corresponds to the initial difference between both algorithms leading to an equal probability for a hybridization event. Thus, in both algorithms the probability for each reaction type is the same.

#### Lemma 2.3.

*The time interval between two successive reactions is the same in both variants*.

*Proof*. Contrary to the Gillespie algorithm where each time increment corresponds to a reaction event, in our new algorithm a time increment is possible without a reaction.

The size of the time step in the Gillespie algorithm is computed by:

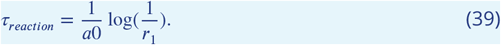

In our algorithm we increment *m* until a reaction happens,i.e., there are *m* − 1 time steps without a reaction. Then the overall time step is:

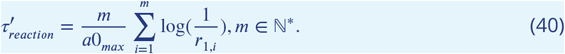

where *r*_1,*i*_ is the *i*-th random number drawn uniformly from [0, 1]. In the following we show that *τ*_*reaction*_ and 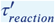 follow the same distribution.

The *m* steps can be seen as sampling with replacement such that either a single reaction with probability *p* = *p*_*reaction*_ takes places or no reaction with probability 1 − *p* takes place. Because replication, hybridization and decay are disjoint events, the probability *p* = *p*_*reaction*_ is the sum of replication, hybridization and decay probabilities:

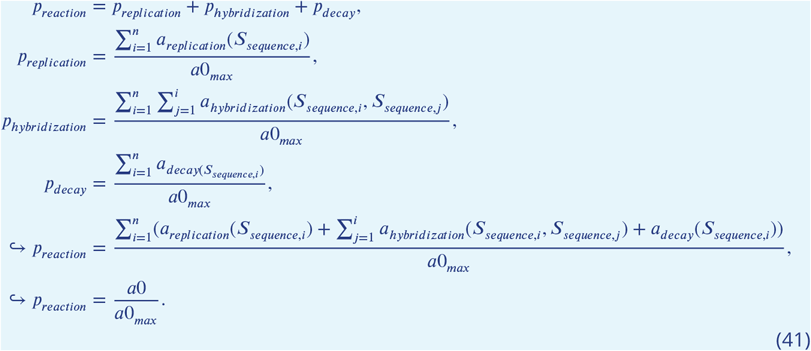

Through sampling with replacement, the number of steps *m* is distributed according to a geometric distribution with expectation 1/*p*. This leads to an expected number of 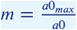 steps. Since a single time step *τ* is distributed according to an exponential distribution with expectation 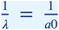 the sum of *m* exponential distributed random variables *X* also has to follow an exponential distribution with the same expectation. As the pdf of such a random variable 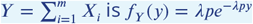 (***Sasha and amWhy, 2014***), *Y* is exponential distributed with expectation 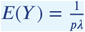.

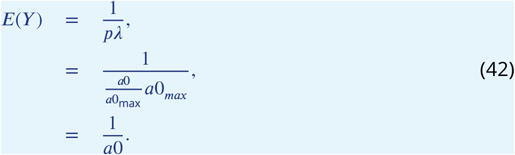

Because *E*(*Y*) corresponds to 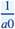, the expectation of the time interval between two reaction events in the Gillespie algorithm, it can be expected that both algorithms have time intervals of equal length between two reactions.

## Appendix 3

### Additional simulation results

#### Validation of accelerated evolution through product inhibition for different target structures

In addition to the standard tRNA shape of length 76 bases, we validated the results using four alternative sequences of length 23, 48, 86, and 91 bases, respectively:

**Appendix 3—table 1.**
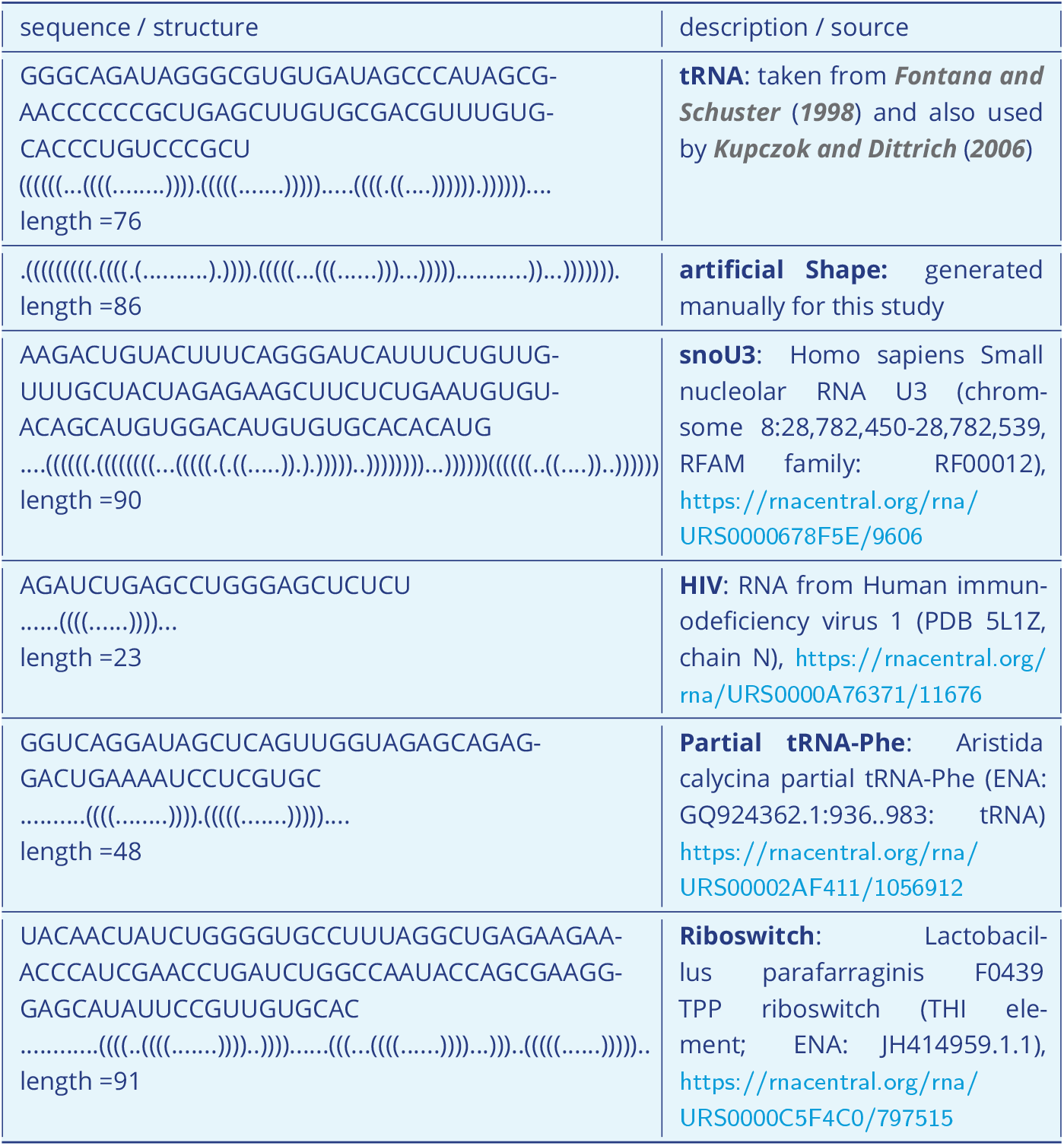
Sequences and structures used in this study. Note that the secondary structures are generated by RNAfold with standard parameters.

**Appendix 3—figure 1.**
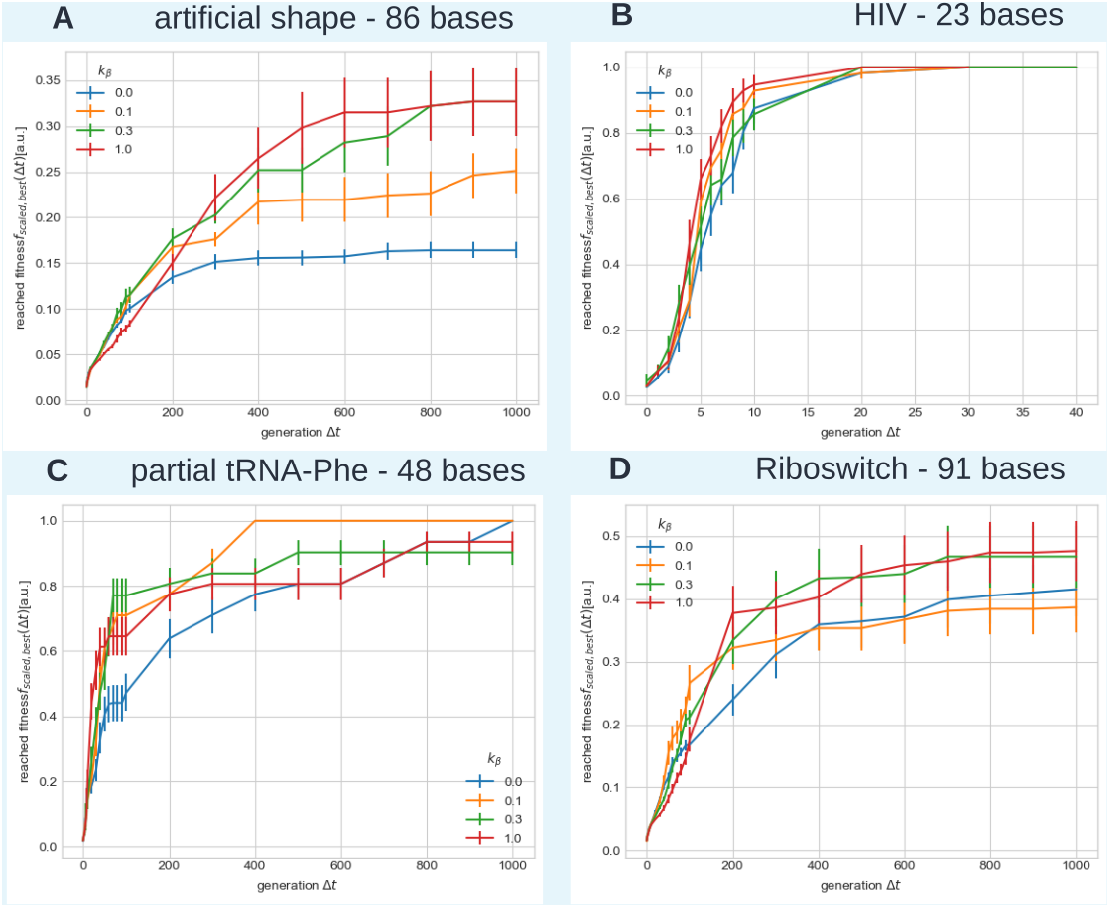
Fitness evolution of the sequence with highest fitness considering (**A**) an artificial shape, (**B**) RNA from Human immunodeficiency virus 1 (PDB 5L1Z, chain N), (**C**) Aristida calycina partial tRNA-Phe, and (**D**) Lactobacillus parafarraginis F0439 TPP riboswitch (THI element). Hybridization rates are *k*_*β*_ = {0, 0.1, 0.3, 1} (blue, orange, green, red).

#### The influence of the hybridization rate on the amount of dimer formation

**Appendix 3—figure 2.**
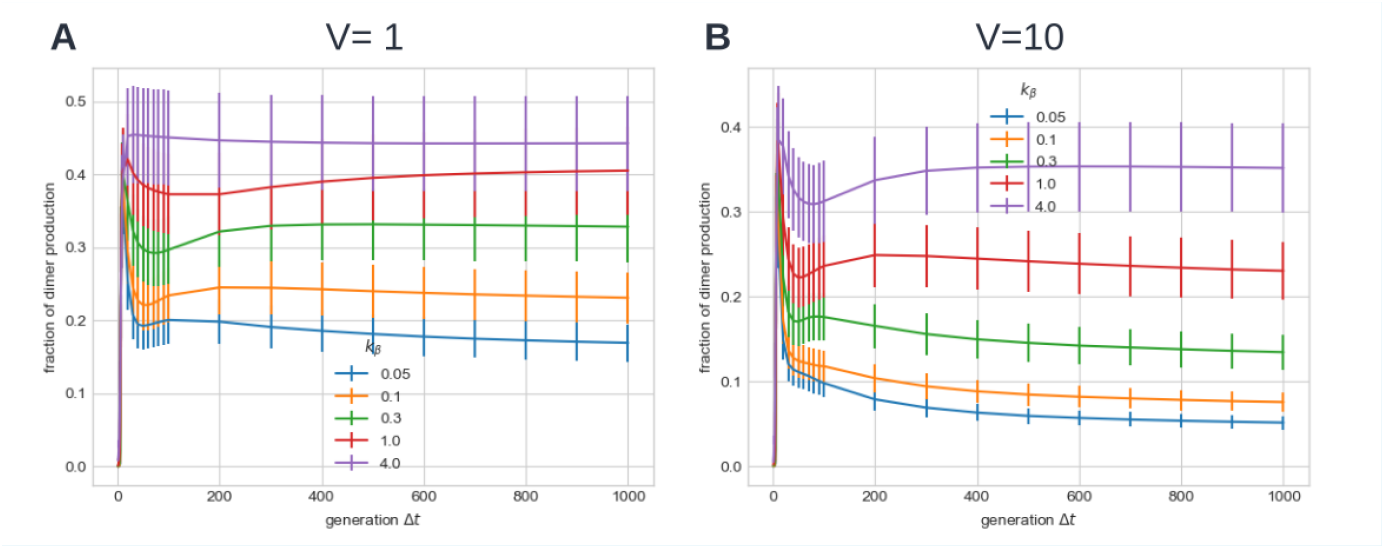
Ratio of the number of hybridization reactions to the number of replication reactions considering a moderate mutation rate *p* = 0.01, (A) a volume *V* = 1, and (B) a volume *V* = 10. The ratio increases with the hybridization strength *k*_*β*_, such that hybridization reactions appear more frequently if *k*_*β*_ is high (here: *k*_*β*_ = 4). An increase in volume by one order of magnitude does not directly correspond to a shift in the fraction of dimer production by one order of magnitude. The depicted ratio is based on the average of 50 simulations, with error bars showing the standard error of the mean, using the standard tRNA of length 76 as the target structure.

#### The observed effects are robust with respect to the chosen hybridization model

**Appendix 3—figure 3.**
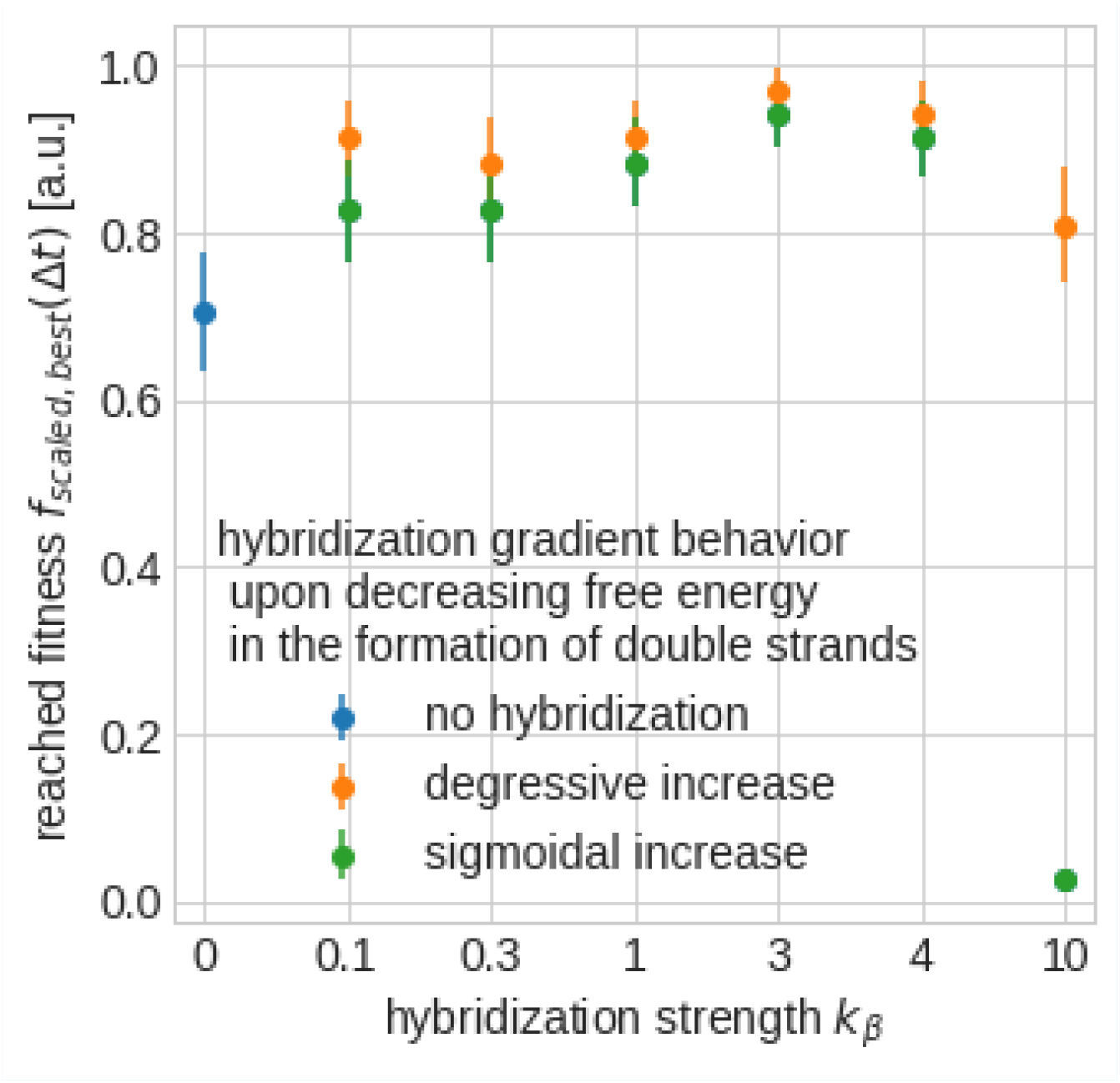
Comparison of two different hybridization models namely the general sigmoidal (green dots) and the degressive (yellow dots) scaling of the hybridization coefficient. Only relatively high (*k*_*β*_ = 10) hybridization rates reveal a significant difference between the two models indicating a greater evolutionary benefit for a degressive increase of the hybridization coefficient. Mean and standard error of the mean shown, based on 25 simulations each, using volume *V* = 10, and a moderate mutation rate *p* = 0.01.

The effect of improved evolution rate through hybridization has been observed using a hybridization coefficient based on the Hamming distance instead of the nearest-neighbor thermodynamics of oligonucleotides (***SantaLucia, 1998***) (data not shown). Furthermore, we varied the scaling. In the results presented so far we applied a sigmoidal scaling for mapping the Gibbs free energy to stochastic rate constants for hybridization (Eq. 8), allowing for a sharp turning between high and low chances for a hybridization event upon colliding sequences. But even for degressive scaling

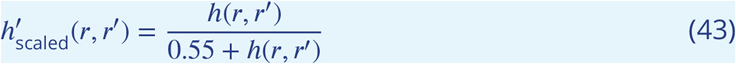

we can observe a similar accelerated evolution (Fig. 3).

## Footnotes

1 The code of the complete model is at https://git.uni-jena.de/ne78xoy/hr-sim-newgillespie.

